# Membrane Insertion of MoS_2_ Nanosheets: Fresh vs. Aged

**DOI:** 10.1101/2020.06.17.158378

**Authors:** Rui Ye, Wei Song, Zonglin Gu, Ruhong Zhou

## Abstract

Fresh two-dimensional (2D) molybdenum disulfide (MoS_2_) can absorb the hydrocarbon contamination from the ambient air and cause surface aging. Thus, understanding how the surface aging process of MoS_2_ affects the interaction with biomolecules is crucial for its applications in the biomedical field. Here, we employed atomistic molecular dynamics simulations to investigate the interactions of fresh and aged MoS_2_ nanosheets with POPE lipid membranes. Our results show that even though both the fresh and aged MoS_2_ nanosheets are capable of spontaneous insertion into the POPE bilayer membrane, the fresh MoS_2_ nanosheet displays significantly more robust interaction than the aged one. The potential mean force (PMF) calculations further confirm that the fresh MoS_2_ nanosheet is more energetically favorable than the aged one in penetrating into the POPE lipid membranes, with the former having ~17 kJ/mol stronger binding affinity than the later. This work provides a deeper understanding of the surface-aging-dependent interaction of MoS_2_ nanosheet with biomolecules, which might help the design of better MoS_2_-based nanodevices with appropriate surface properties.

## Introduction

Molybdenum disulfide (MoS_2_) is a member of the so-called transition-metal dichalcogenide (TMD) family, which has received great attention lately due to their unique physical and chemical properties. Previous studies have shown wide applications of MoS_2_ nanosheets in optoelectronics,^1^ field emission transistors,^2^ gas sensors^3–4^ and hydrogen storage.^5^ More recently, MoS_2_ nanosheets have also shown increasing interest in the biomedical field. For example, it has been reported that their strong near-infrared (NIR) absorption feature enables them to be used for photothermal therapy in cancer treatment.^6–7^ Also, the moderate direct bandgap of MoS_2_ nanosheet has been explored for protein and DNA detections.^8–9^ While the strong absorbance of X-ray by Mo atoms makes it an appealing contrast agent in X-ray computed tomography imaging.^7^

Recent studies also demonstrate that the functionalized MoS_2_ nanosheets have high inhibitory and bactericidal activities against ESKAPE pathogens by destructing their cell membranes.^10^ On the other hand, Yang *et al.* proposed that the antibacterial effects of MoS_2_ nanosheets to the *Escherichia coli* (*E. coli*) is attributed by the chemical oxidation and membrane stress mechanisms.^11^ In addition, MoS_2_ nanosheets can also disrupt the integrity of the cell membranes and extract the phospholipids from the membrane bilayers.^12^ These investigations have shown that the MoS_2_ nanosheets hold strong interactions with cell membranes and other biomolecules, which makes them appealing to studies of nanotoxicity (and potential nanomedicine as antibacterial agents).

It is worth noting that the freshly prepared MoS_2_ nanosheets (fresh MoS_2_) can absorb hydrocarbons when they are exposed to air and induce surface aging (named as aged MoS_2_), which thereby affects the surface hydrophobicity and topology and causes changes in the water contact angles (WCA) on the MoS_2_ surface as evidenced by a series of variable values measured by different experiments.^13–16^ More importantly, the surface aging process of a MoS_2_ nanosheet may result in a potential effect on its interaction with biomolecules at the bio-nano interface, which is yet unknown to a large extent.

In this paper, we employ molecular dynamics (MD) simulations to investigate the underlying mechanism of the interactions between the MoS_2_ nanosheets (with fresh vs. aged) and the POPE lipid membrane, aiming to understand how the surface aging process of MoS_2_ affects interactions with biomolecules. Based on MD simulations, together with free energy calculations, we found that the fresh MoS_2_ nanosheet presents a stronger binding affinity to the POPE lipid membranes than the aged one. The results suggest that the surface aging process of MoS_2_ could affect the interaction with biomolecules.

## Results and Discussion

Here, we applied the MD simulations to investigate the interactions of fresh and aged MoS_2_ nanosheets with POPE lipid membrane. In order to efficiently model both fresh and aged MoS_2_ surfaces, we have developed a simplified model by only adjusting the Lennard-Jones parameter, ε_S_, (the depth of the potential well of a sulfur atom)^17^, which demonstrates a surprisingly linear relationship between the ε_S_ and water contact angle (WCA). In this study, the surface-aging dependent interaction of a MoS_2_ nanosheet with membranes can thus be “achieved” using this simple MoS_2_ force field model.^17–18^ **Table 1** lists two sets of force field parameters for the fresh and aged MoS_2_ nanosheets, respectively.

**Table 1.**
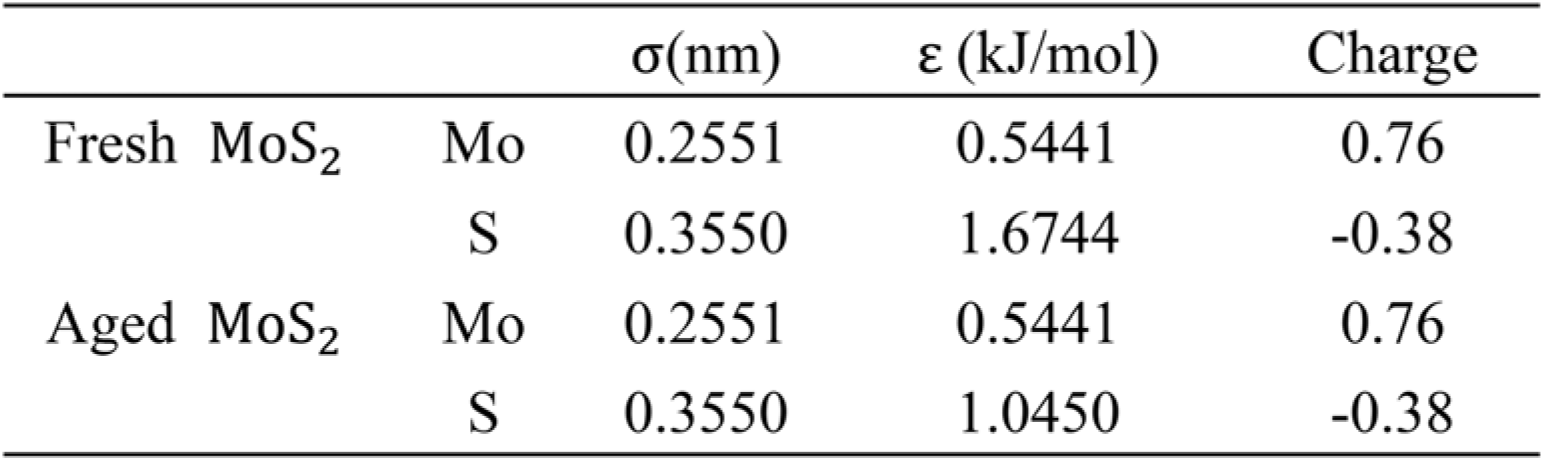
Force field parameters for the fresh and aged MoS_2_ nanosheets, respectively. (Derived from our previous work ^17-18^)

The initial configuration of a MoS_2_ nanosheet and POPE lipid membrane is illustrated in **Fig. 1** (details can be seen in the Methods section). **Fig. 2** shows the final conformations of the fresh and aged MoS_2_ nanosheets inserting into the lipid membrane from three independent trajectories. As can be clearly seen from **Fig. 2** that both the fresh and aged MoS_2_ nanosheets finally bury themselves into the lipid membranes with their orientations perpendicular to the membrane surfaces. In addition, both MoS_2_ nanosheets (with fresh vs. aged) are mostly in direct contact with the hydrophobic tail of the lipid molecules and parallel to the tail chains. The results reveal that both the fresh and aged MoS_2_ nanosheets can penetrate into the POPE lipid bilayer membrane, independent of their surface properties.

**Figure 1.**
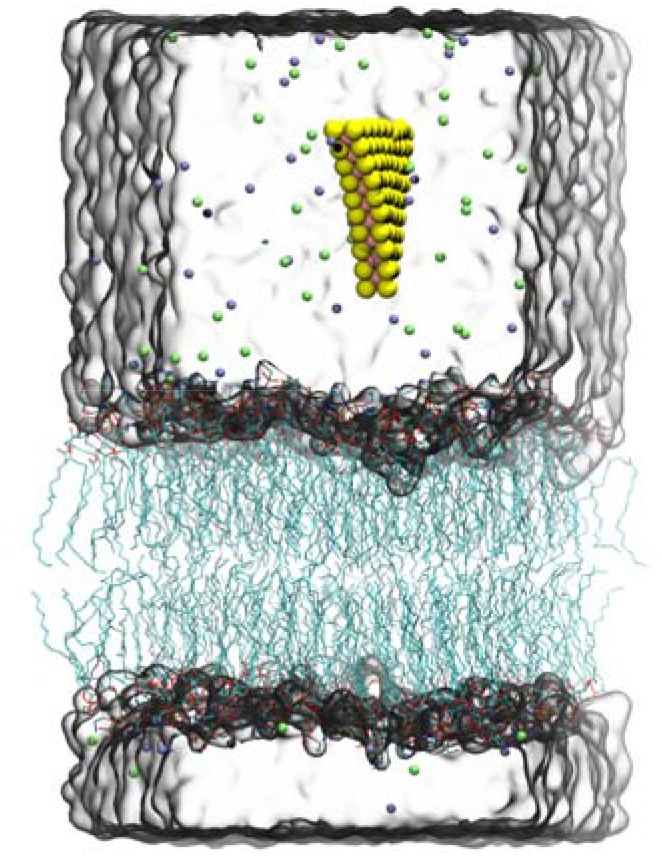
The simulation model of MoS_2_ nanosheet on the POPE lipid membrane in a water box. Molybdenum and sulfur atoms are displayed by pink and yellow spheres, respectively. The phospholipids are represented in blue lines with P atoms as red lines. Sodium and chlorine ions are shown as green and purple spheres, respectively.

**Figure 2.**
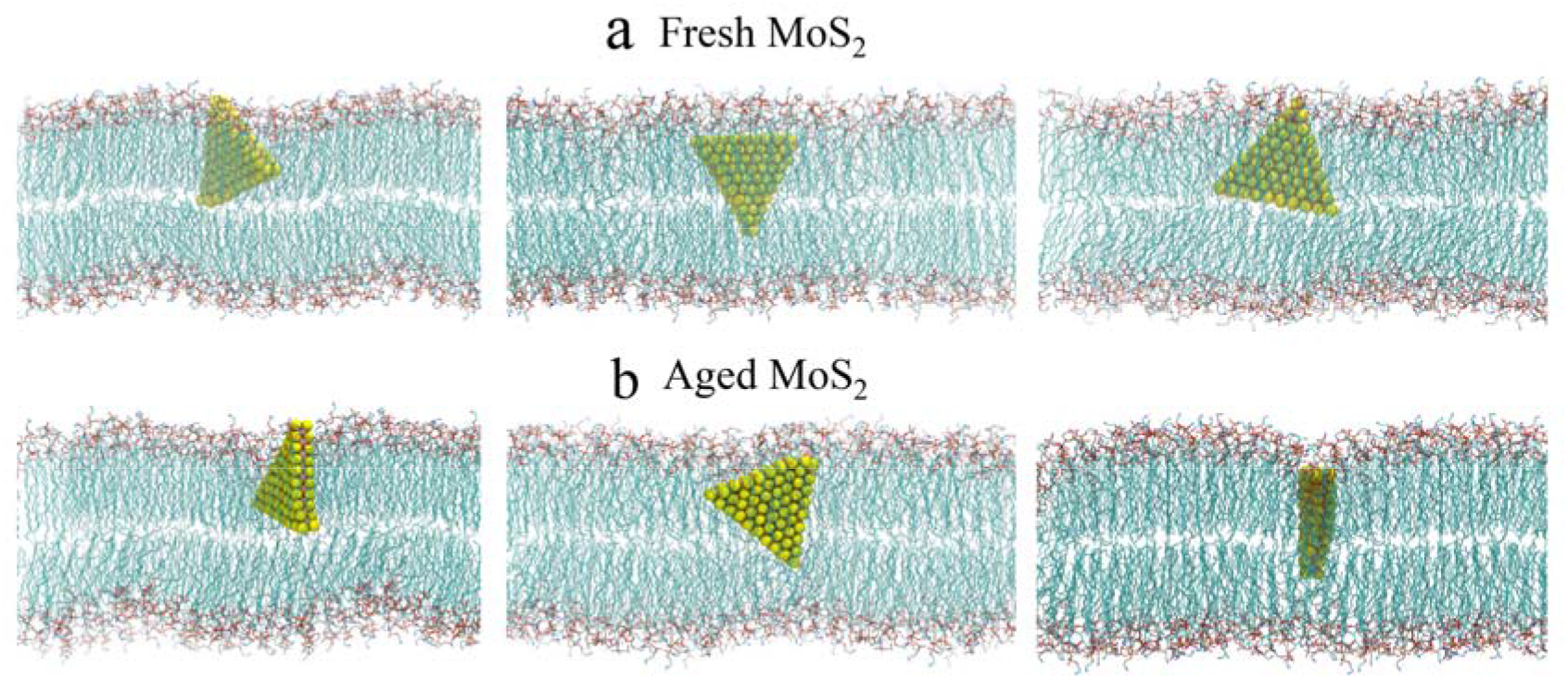
Last snapshots of the fresh MoS_2_ nanosheet (a) and aged MoS_2_ nanosheet (b) on the POPE lipid membranes from three independent trajectories at 200 ns.

To better understand the interaction process, we further analyzed the time evolution of the atom contact number between the MoS_2_ nanosheets (fresh vs. aged) and lipid membrane (**Fig. 3a and c**). Here, the atom contact number was recorded when the distance between a MoS_2_ nanosheet and any heavy atom of lipid molecules is less than 0.5 nm. In addition, the binding conformations of the fresh and aged MoS_2_ with the POPE membrane at four key snapshots were also illustrated in **Fig. 3b and d**. Initially, in the case of the fresh MoS_2_ nanosheet, it moved freely in the water and has intermittent contacts with the lipid membrane, yielding an atom contact number less than 20 (0-5 ns). At t = 5 ns, fresh MoS_2_ started to contact the membrane with a point-to-face orientation pointing to the target membrane. At t = 18 ns, the fresh MoS_2_ has changed its binding conformation into a face-to-face orientation with an increase in atom contact number close to 100. At t = 22 ns, the fresh MoS_2_ nanosheet has changed its conformation and is to tilt itself with one of the vertexes inserting into the membrane, along with a transient decrease in contact number compared to the value at 18 ns. Starting from t = 22 ns, the fresh MoS_2_ continued to penetrate into the membrane with a dramatic increase in the atom contact number. After 65 ns, the interactions between the fresh MoS_2_ and POPE membrane reached an equilibrium state with limited fluctuation in contact number (~275). As for the aged MoS_2_, the general trend in the change of atom contact numbers and binding conformations is similar to the aforementioned fresh nanosheet as evidenced by the illustrations in **Fig. 3c** and **d**. These results suggest that both the MoS_2_ nanosheets (fresh vs. aged) are capable of penetrating into the POPE lipid membranes.

**Figure 3.**
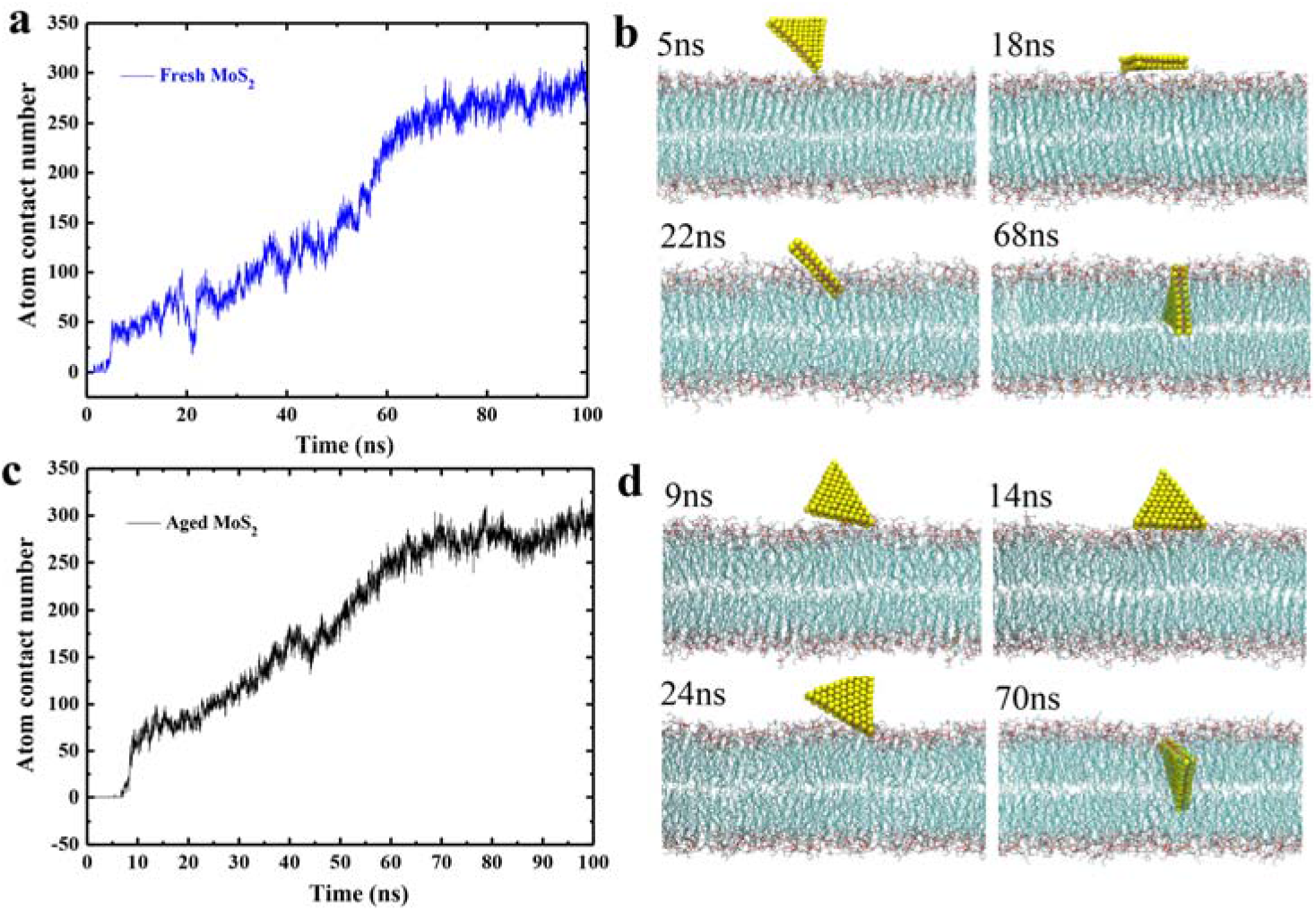
The number of contact between the fresh (a)/aged (c) MoS_2_ nanosheets and POPE lipid membranes along 100 ns simulations. The binding conformations of the fresh (b) and aged (d) MoS_2_ nanosheets to POPE lipid membranes at four key time points.

We then analyzed the time evolution of the interaction energy between the MoS_2_ nanosheets (fresh vs. aged) and lipid membrane to further illustrate the insertion processes. As can be seen from **Fig 4a** and **b**, the van der Waals (vdW) force dominates the interaction energies in both fresh and aged MoS_2_ nanosheets with POPE lipid membrane. Moreover, the vdW interaction energy of fresh MoS_2_ is lower than that of aged MoS_2_ in interacting with the POPE lipid membrane, whereas the Coulombic (Coul) interaction energy of fresh MoS_2_ is slightly higher than that of aged MoS_2_ (**Fig. S1**). Further analysis showed that the atom contact number between the fresh MoS_2_ and head group is lower than that between the aged MoS_2_ and head group. Conversely, the opposite trend is observed for the fresh and aged MoS_2_ with the tail groups (**Fig. 4c**). In addition, **Fig. 4d** showed that the distance between the center of mass (COM) of the fresh MoS_2_ and lipid membrane is shorter than that of the aged MoS_2_ with the membrane, indicating that the fresh MoS_2_ might have more damage to the cell membrane.

**Figure 4.**
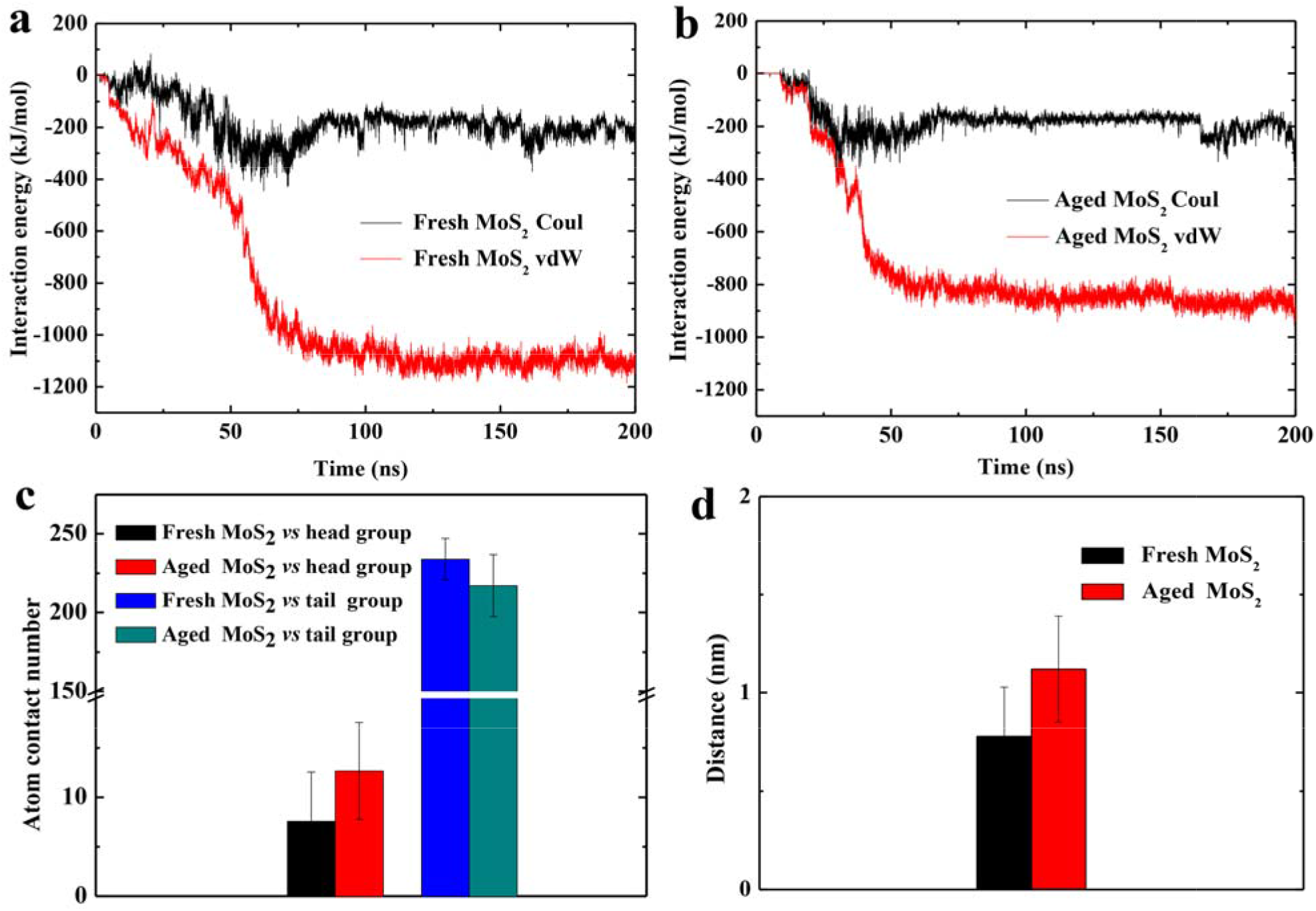
The interaction energies (including vdW and Coul energies) between the fresh (a) and aged (b) MoS_2_ nanosheets and lipid membrane during simulations. (c) The number of contact between fresh/aged MoS_2_ nanosheets and head/tail groups of lipid membranes. (d) The distance between the center of mass (COM) of the MoS_2_ nanosheets (fresh vs. aged) and lipid membranes at equilibrium.

Additionally, we also calculated the binding free energies between the MoS_2_ nanosheets (fresh vs. aged) and POPE lipid membranes by monitoring the relative free energy when pulling the MoS_2_ nanosheets along the Z-direction perpendicular to the membrane, using the potential of mean force (PMF) method (details can be found in the Methods section). As seen in **Fig. 5** that the free energy of the fresh MoS_2_ is ~17 kJ/mol stronger than the aged one, which indicates that the fresh MoS_2_ nanosheet is more energetically favorable to penetrate into the cell membrane than the aged one, though both the fresh and aged MoS_2_ nanosheets exhibit spontaneous penetrating behaviors to the POPE membrane. Besides, the minimum of the PMF curves corresponds to the COM distance of 0.8 nm and 1.1 nm for the fresh and aged MoS_2_, respectively, in agreement with **Fig. 4d** and consistent with the indication that the fresh MoS_2_ might have more damage to the cell membrane.

**Figure 5.**
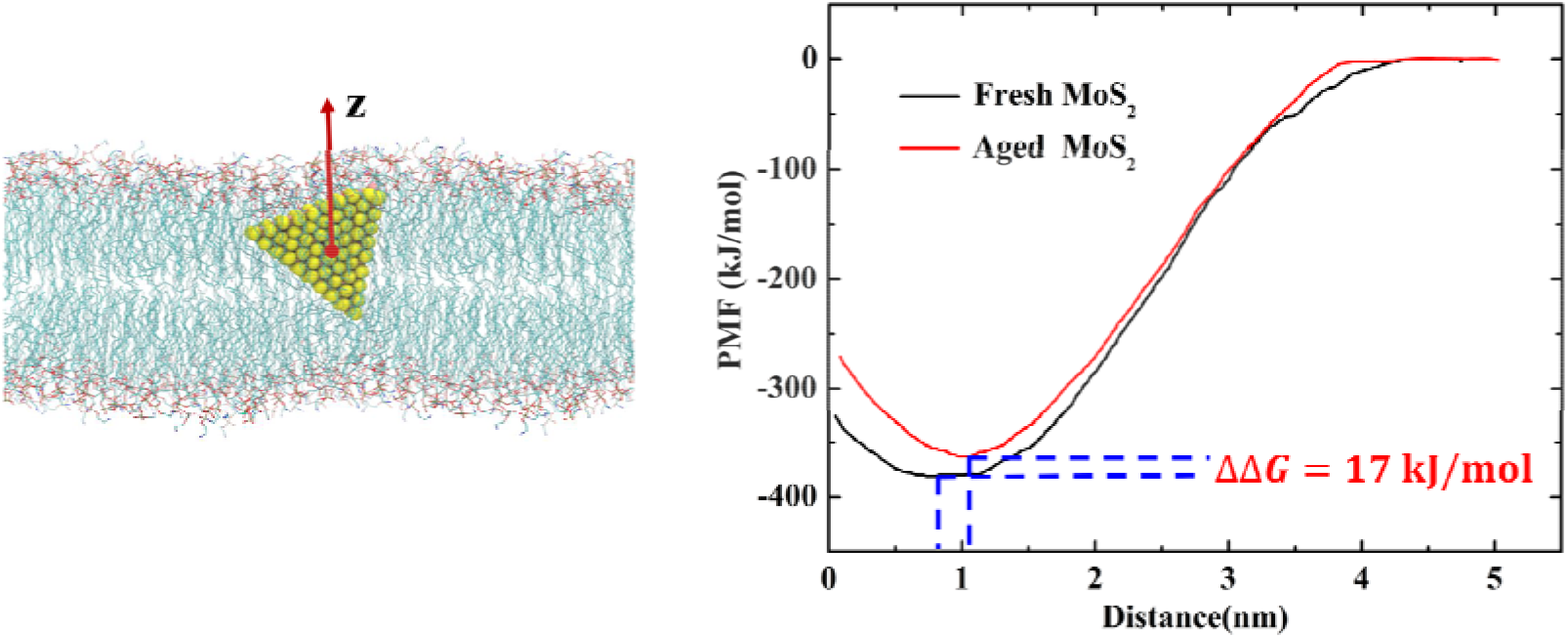
Potential of mean force (PMF) curve showing the free energy as the MoS_2_ nanosheet moving along the Z-axis direction, normal to the membrane surface.

## Conclusion

In this work, we employed atomistic molecular dynamics simulations to investigate the interactions between MoS_2_ nanosheets (with fresh vs. aged) and the POPE lipid membrane, aiming to understand how the surface aging process of a MoS_2_ nanosheet affects its interaction with cell membranes. Our results show that even though both the fresh and aged MoS_2_ nanosheets are able to insert into POPE bilayers, the fresh MoS_2_ nanosheet presents significantly more robust interaction with the membrane than the aged one. Free energy calculations computed by the potential mean force (PMF) further verify that the membrane insertion process of the fresh MoS_2_ nanosheet is more energetically favorable than the aged one, with the former showing ~17 kJ/mol more binding affinity than the latter. Our findings reveal that MoS_2_ has a surface-aging-dependent interaction with cell membranes, which may be beneficial to future applications in biomedicine.

## Methods

### Molecular dynamics simulation

In this work, the triangular MoS_2_ nanosheet (side length of 2.89 nm) was used due to its small size and was also proved to have the capacity in penetrating into the bilayer.^19^ The membrane with surface dimensions of 6.25 × 6.25 nm^2^ was constructed by using CHARMM-Gui (http://www.charmm-gui.org), yielding 168 palmitoyloleoylphosphatidylethanolamine (POPE) lipids.^20^ Two types of MoS_2_ nanosheets (with fresh vs. aged) were placed above the lipid membrane with a minimum distance of about 1.2 nm (as shown in **Fig. 1**). Two systems were placed in the same box (6.25 × 6.25 × 12.73 nm), containing 10473 water molecules, 48 Na ions and 48 Cl ions.

All MD simulations are carried out with the software package GROMACS (version 5.1.4).^21^ VMD software was utilized to visualize MD trajectories and draw snapshots.^22^ The force field parameters of the fresh and aged MoS_2_ nanosheets were employed based on our previous works.^17–18^ The CHARMM 36 force field ^23–25^ and TIP3P water model ^26^ were adopted for POPE molecules and water molecules, respectively. Following similar protocols in our previous studies^27–33^, periodic boundary conditions of these systems were treated in all directions (x y and z). The temperature and pressure were fixed at 300 K and 1 atm using v-rescale thermostat^34^ and semi-isotropic Parrinello-Rahman algorithm (x+y, z).^35^ The long-range electrostatic interactions were treated with the PME method,^36^ and the van der Waals (vdW) interactions were calculated with a cutoff distance of 1.2 nm. The geometrical properties of solute bonds associated with hydrogen were kept constant at their equilibrium values with the LINCS algorithm,^37^ and water geometry was also constrained using the SETTLE algorithm.^38^ All simulations were performed for 200 ns and data were collected every 10 ps.

### Potential of mean force (PMF)

The PMF profiles of the fresh and aged MoS_2_ nanosheets along the z-direction perpendicular to the membrane surface were calculated using umbrella sampling simulations.^39–41^ The distance (d) to the center of the membrane was restrained at a reference distance (d_0_) with a harmonic force

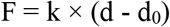

where k is the force constant (2000 kJ mol^-1^ nm^-2^). The spacing of the sampling windows was 0.1 nm. For each d_0_, the system was equilibrated for 2 ns, followed by a 10-ns productive run. The free energy profiles were obtained by the Weighted Histogram Analysis Method.^42–44^

## ACKNOWLEDGEMENT

We thank Mei Feng and Jianxiang Huang for help with the manuscript. This work is partially supported by the National Natural Science Foundation of China (Grants 11574224 and U1967217) and China Postdoctoral Science Foundation (Grant 2019M652069 and 2019T120506). R.Z. also acknowledges the financial support from W. M. Keck Foundation (Grant award 2019-2022).

## Reference

1. Splendiani, A.; Sun, L.; Zhang, Y.; Li, T.; Kim, J.; Chim, C. Y.; Galli, G.; Wang, F., Emerging photoluminescence in monolayer mos2. Nano Lett. 2010, 10 (4), 1271–1275.

2. Radisavljevic, B.; Radenovic, A.; Brivio, J.; Giacometti, V.; Kis, A., Single-layer mos2 transistors. Nat Nanotechnol 2011, 6 (3), 147–150.

3. Qiu, H.; Pan, L.; Yao, Z.; Li, J.; Shi, Y.; Wang, X. J. A. P. L., Electrical characterization of back-gated bi-layer mos2 field-effect transistors and the effect of ambient on their performances. Appl. Phys. Lett. 2012, 100 (12), 123104.

4. Perkins, F. K.; Friedman, A. L.; Cobas, E.; Campbell, P. M.; Jernigan, G. G.; Jonker, B. T., Chemical vapor sensing with monolayer mos2. Nano Lett. 2013, 13 (2), 668–673.

5. Chen, J.; Kuriyama, N.; Yuan, H.; Takeshita, H. T.; Sakai, T. J. J. o. t. A. C. S., Electrochemical hydrogen storage in mos2 nanotubes. J. Am.Chem. Soc. 2001, 123 (47), 11813–11814.

6. Wang, S.; Li, K.; Chen, Y.; Chen, H.; Ma, M.; Feng, J.; Zhao, Q.; Shi, J., Biocompatible pegylated mos2 nanosheets: Controllable bottom-up synthesis and highly efficient photothermal regression of tumor. Biomaterials 2015, 39, 206–217.

7. Yin, W.; Yan, L.; Yu, J.; Tian, G.; Zhou, L.; Zheng, X.; Zhang, X.; Yong, Y.; Li, J.; Gu, Z., et al., High-throughput synthesis of single-layer mos2 nanosheets as a near-infrared photothermal-triggered drug delivery for effective cancer therapy. ACS Nano 2014, 8 (7), 6922–6933.

8. Wang, L.; Wang, Y.; Wong, J. I.; Palacios, T.; Kong, J.; Yang, H. Y., Functionalized mos2 nanosheet-based field-effect biosensor for label-free sensitive detection of cancer marker proteins in solution. Small 2014, 10 (6), 1101–1105.

9. Zhu, C. F.; Zeng, Z. Y.; Li, H.; Li, F.; Fan, C. H.; Zhang, H., Single-layer mos2-based nanoprobes for homogeneous detection of biomolecules. J Am Chem Soc 2013, 135 (16), 5998–6001.

10. Pandit, S.; Karunakaran, S.; Boda, S. K.; Basu, B.; De, M., High antibacterial activity of functionalized chemically exfoliated mos2. Acs Appl Mater Inter 2016, 8 (46), 31567–31573.

11. Yang, X.; Li, J.; Liang, T.; Ma, C. Y.; Zhang, Y. Y.; Chen, H. Z.; Hanagata, N.; Su, H. X.; Xu, M. S., Antibacterial activity of two-dimensional mos_2_ sheets. Nanoscale 2014, 6 (17), 10126–10133.

12. Wu, R. R.; Ou, X. W.; Tian, R. R.; Zhang, J.; Jin, H. S.; Dong, M. D.; Li, J. Y.; Liu, L., Membrane destruction and phospholipid extraction by using two-dimensional mos2 nanosheets. Nanoscale 2018, 10 (43), 20162–20170.

13. Kelebek, S., Critical surface-tension of wetting and of floatability of molybdenite and sulfur. J Colloid Interf Sci 1988, 124 (2), 504–514.

14. Zhang, S.; Zeng, X. T.; Tang, Z. G.; Tan, M. J., Exploring the antisticking properties of solid lubricant thin films in transfer molding. Int J Mod Phys B 2002, 16 (6-7), 1080–1085.

15. Gaur, A. P. S.; Sahoo, S.; Ahmadi, M.; Dash, S. P.; Guinel, M. J. F.; Katiyar, R. S., Surface energy engineering for tunable wettability through controlled synthesis of mos2. Nano Lett. 2014, 14 (8), 4314–4321.

16. Kozbial, A.; Gong, X.; Liu, H. T.; Li, L., Understanding the intrinsic water wettability of molybdenum disulfide (mos2). Langmuir 2015, 31 (30), 8429–8435.

17. Zhang, L.; Luan, B.; Zhou, R., Parameterization of molybdenum disulfide interacting with water using the free energy perturbation method. J. Phys. Chem. B 2019, 123 (34), 7243–7252.

18. Luan, B.; Zhou, R., Wettability and friction of water on a mos2 nanosheet. Appl. Phys. Lett. 2016, 108 (13), 131601.

19. Gu, Z.; Chen, S. H.; Ding, Z.; Song, W.; Wei, W.; Liu, S.; Ma, G.; Zhou, R., The molecular mechanism of robust macrophage immune responses induced by pegylated molybdenum disulfide. Nanoscale 2019, 11 (46), 22293–22304.

20. Jo, S.; Lim, J. B.; Klauda, J. B.; Im, W., Charmm-gui membrane builder for mixed bilayers and its application to yeast membranes. Biophys. J. 2009, 97 (1), 50–58.

21. Hess, B.; Kutzner, C.; van der Spoel, D.; Lindahl, E., Gromacs 4: Algorithms for highly efficient, load-balanced, and scalable molecular simulation. J. Chem. Theory Comput. 2008, 4 (3), 435–447.

22. Humphrey, W.; Dalke, A.; Schulten, K., Vmd: Visual molecular dynamics. J Mol Graph Model 1996, 14 (1), 33–38.

23. Brooks, B. R.; Bruccoleri, R. E.; Olafson, B. D.; States, D. J.; Swaminathan, S.; Karplus, M., Charmm - a program for macromolecular energy, minimization, and dynamics calculations. J Comput Chem 1983, 4 (2), 187–217.

24. Klauda, J. B.; Venable, R. M.; Freites, J. A.; O’Connor, J. W.; Tobias, D. J.; Mondragon-Ramirez, C.; Vorobyov, I.; MacKerell, A. D.; Pastor, R. W., Update of the charmm all-atom additive force field for lipids: Validation on six lipid types. J. Phys. Chem. B 2010, 114 (23), 7830–7843.

25. Mackerell, A. D.; Feig, M.; Brooks, C. L., Extending the treatment of backbone energetics in protein force fields: Limitations of gas-phase quantum mechanics in reproducing protein conformational distributions in molecular dynamics simulations. J Comput Chem 2004, 25 (11), 1400–1415.

26. Jorgensen, W. L.; Chandrasekhar, J.; Madura, J. D.; Impey, R. W.; Klein, M. L., Comparison of simple potential functions for simulating liquid water. J Chem Phys 1983, 79 (2), 926–935.

27. Ahmed, R.; Omidian, Z.; Giwa, A.; Cornwell, B.; Majety, N.; Bell, D. R.; Lee, S.; Zhang, H.; Michels, A.; Desiderio, S., et al., A public bcr present in a unique dual-receptor-expressing lymphocyte from type 1 diabetes patients encodes a potent t cell autoantigen. Cell 2019, 177 (6), 1583–1599 e16.

28. Luo, N.; Weber, J. K.; Wang, S.; Luan, B.; Yue, H.; Xi, X.; Du, J.; Yang, Z.; Wei, W.; Zhou, R., et al., Pegylated graphene oxide elicits strong immunological responses despite surface passivation. Nat Commun 2017, 8, 14537.

29. Fitch, B. G.; Rayshubskiy, A.; Eleftheriou, M.; J., C. T.; Giampaga, M.; Zhestkov, Y.; Pitman, M. C.; Suits, F.; Grossfield, A.; Pitera, J., et al., Blue matter: Strong scaling of molecular dynamics on blue gene/l. Springe Berlin Heidelberg: International Conference on Computational Science, 2006.

30. Kaminski, G. A.; Friesner, R. A.; Zhou, R., A computationally inexpensive modification of the point dipole electrostatic polarization model for molecular simulations. J. Comput. Chem. 2003, 24 (3), 267–76.

31. Li, J.; Liu, T.; Li, X.; Ye, L.; Chen, H.; Fang, H.; Wu, Z.; Zhou, R., Hydration and dewetting near graphite-ch(3) and graphite-cooh plates. J Phys Chem B 2005, 109 (28), 13639–48.

32. Chowell, D.; Morris, L. G. T.; Grigg, C. M.; Weber, J. K.; Samstein, R. M.; Makarov, V.; Kuo, F.; Kendall, S. M.; Requena, D.; Riaz, N., et al., Patient hla class i genotype influences cancer response to checkpoint blockade immunotherapy. Science 2018, 359 (6375), 582–587.

33. Zhou, R., Exploring the protein folding free energy landscape: Coupling replica exchange method with p3me/respa algorithm. J. Mol. Graph. Model. 2004, 22 (5), 451–63.

34. Bussi, G.; Donadio, D.; Parrinello, M., Canonical sampling through velocity rescaling. J. Chem. Phys. 2007, 126 (1), 014101.

35. Parrinello, M.; Rahman, A., Polymorphic transitions in single-crystals - a new molecular-dynamics method. J Appl Phys 1981, 52 (12), 7182–7190.

36. Darden, T.; York, D.; Pedersen, L., Particle mesh ewald - an n.Log(n) method for ewald sums in large systems. J. Chem. Phys. 1993, 98 (12), 10089–10092.

37. Hess, B.; Bekker, H.; Berendsen, H. J. C.; Fraaije, J. G. E. M., Lincs: A linear constraint solver for molecular simulations. J Comput Chem 1997, 18 (12), 1463–1472.

38. Miyamoto, S.; Kollman, P. A., Settle - an analytical version of the shake and rattle algorithm for rigid water models. J Comput Chem 1992, 13 (8), 952–962.

39. Kumar, S.; Rosenberg, J. M.; Bouzida, D.; Swendsen, R. H.; Kollman, P. A., Multidimensional free-energy calculations using the weighted histogram analysis method. J Comput Chem 1995, 16 (11), 1339–1350.

40. Roux, B., The calculation of the potential of mean force using computer-simulations. Comput Phys Commun 1995, 91 (1-3), 275–282.

41. Torrie, G. M.; Valleau, J. P., Non-physical sampling distributions in monte-carlo free-energy estimation - umbrella sampling. J Comput Phys 1977, 23 (2), 187–199.

42. Efron, B., 1977 rietz lecture - bootstrap methods - another look at the jackknife. Ann Stat 1979, 7 (1), 1–26.

43. Hub, J. S.; de Groot, B. L.; van der Spoel, D., G_wham-a free weighted histogram analysis implementation including robust error and autocorrelation estimates. J. Chem. Theory Comput. 2010, 6 (12), 3713–3720.

44. Kirkwood, J. G., Statistical mechanics of fluid mixtures. J. Chem. Phys. 1935, 3 (5), 300–313.

