## supplementary figure S1 for "Membrane Insertion of MoS_2_ Nanosheets: Fresh vs. Aged"

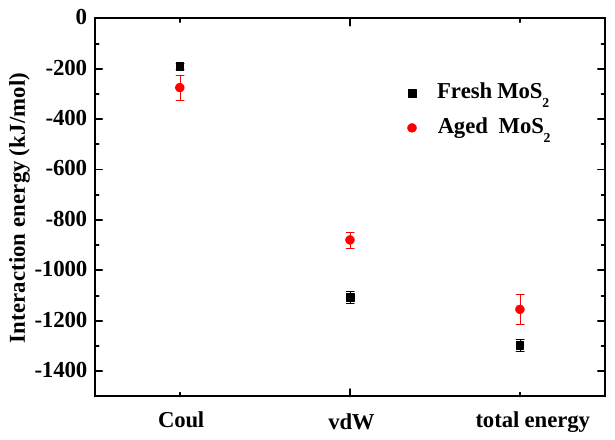

**Figure S1.** Coulombic (Coul), van der Waals (vdW) and total energies averaged from the last 10ns of the simulations out of three parallel trajectories.
